# A New Software Tool for Computer Assisted In Vivo High-Content Analysis of Transplanted Fluorescent Cells in intact Zebrafish Larvae

**DOI:** 10.1101/2022.07.02.498535

**Authors:** Jan-Lukas Førde, Ingeborg Nerbø Reiten, Kari Espolin Fladmark, Astrid Olsnes Kittang, Lars Herfindal

## Abstract

Acute myeloid leukaemia and myelodysplastic syndromes are cancers of the bone marrow with poor prognosis in frail and older patients. To investigate cancer pathophysiology and therapies, confocal imaging of fluorescent cancer cells, and their response to treatments in zebrafish larvae, yields valuable information. While zebrafish larvae are well suited for confocal imaging, the lack of efficient processing of large datasets remains a severe bottleneck. To alleviate this problem, we present a software-tool to segment cells from confocal images and track characteristics such as volume, location in the larva and fluorescent intensity on a single-cell basis. Using this software-tool, we were able to characterize the responses of the cancer cell lines Molm-13 and MDS-L to established treatments. By utilizing the computer-assisted processing of confocal images as presented here, more information can be obtained while being less time-consuming and reducing the demand of manual data handling, when compared to a manual approach, thereby accelerating the pursuit of novel anti-cancer treatments.

**Summary statement:** We present a software tool for automatic cell segmentation of fluorescent cancer cells in zebrafish larvae to determine characteristics of the cancer cell population on a single-cell basis.

## Introduction

The myeloid malignancies acute myeloid leukaemia (AML) and myelodysplastic syndromes (MDS) are related cancers of the bone marrow, both of which are highly. Common for both diseases are their clinical manifestations with cytopenia and bone marrow failure (Cazzola, 2020; Döhner et al., 2015; Mohammad, 2018). In approximately 20% of MDS patients, the disease can progress to secondary AML, which is associated with poor prognosis and low survival (Menssen and Walter, 2020). AML and MDS are rare in children and young adults, and both diseases have a median age of diagnosis of 70 (https://www.cancer.org/cancer/acute-myeloid-leukemia/about/key-statistics.html and https://www.cancer.org/cancer/myelodysplastic-syndrome/about/key-statistics.html retrieved 4/28/2022). While alternative treatments such as hematopoietic stem cell transplantation (HTC) and immunotherapies like CAR-t-cell therapy exist, most patients rely on chemotherapy. For AML, the standard induction therapy consists of continuously infused cytarabine for seven days and daily administration of an anthracycline such as daunorubicin for three (Döhner et al., 2015). While MDS patients in a low-risk group can be treated symptomatically with blood transfusions, high-risk patients are treated with hypomethylating agents like azacitidine (Aza), or in younger patients exhibiting high blast counts with chemotherapy as in AML (Encyclopedia of Molecular Mechanisms of Disease, 2009). A challenge in the treatment of MDS and AML is that elderly or patients with poor general condition are unsuited for curative treatments like HTC, and have to resort to chemotherapy (Saygin and Carraway, 2021). Moreover, these patients also have low tolerance for intensive chemotherapy. For AML, the median survival is 5 to 10 months in elderly and frail patients, and for high-risk MDS patients, the only remaining treatment is hypomethylating agents like Aza, which gives an overall median survival of 10 months compared to supportive care (Döhner et al., 2015; Fenaux et al., 2009). While some new treatments are in clinical trials or have been approved in the last decade, such as Enasidenib for AML and MDS patients with IDH2 mutation, a demand for novel treatments persist (Cazzola, 2020). Especially considering the heterogeneity of both AML and MDS, new disease models are needed to facilitate the rapid development of novel anti-cancer drugs.

Zebrafish have become an intriguing tool for the development of anti-cancer therapies. The species shares around 70% of human genes and has orthologues for approximately 80% of proteins linked to human diseases (Molina et al., 2021). Compared to mammalian disease models like mice, zebrafish possess several benefits such as a high fecundity rate, low maintenance cost and ease of genetic manipulation. Zebrafish are also translucent during the early stages of development or even throughout adulthood in the genetically modified zebrafish line Casper, which enables the use of optical microscopy techniques on living subjects (White et al., 2008). Especially well-suited for drug screening are zebrafish during the embryonal and larval period, due to their aforementioned optical transparency, small size, and absence of an adaptive immune system until four to six weeks post fertilization, while still having important anatomical structures and physiological processes (Molina et al., 2021; Sullivan et al., 2017).

In cancer research, observing the response of cancer cells to treatments, as well as interactions between the cancer and host, is highly valuable. By injecting fluorescently labelled cancer cells in zebrafish embryo and larvae, these interactions can be observed using confocal microscopy. While zebrafish embryos and larvae are well suited for microscopy, processing of the acquired data remains a significant bottleneck (Mikut et al., 2013). Automated processing methods can enable larger scale high-content screenings using confocal microscopy where the manual data processing and analysis limits the amount of information that can be mined. Some solutions of this problem are already devised (Carreira et al., 2021; Yamamoto et al., 2019); however, these solutions are optimized to work on two-dimensional microscope images. This approach struggles to identify individual cells if these are overlapping in the image plane, hence it identifies fluorescent areas instead of individual cells. This approach has some shortfalls, for instance inaccurate determination of cancer cell locations and colocalization with other fluorescent elements of interests due to the lack of one spatial coordinate. Additionally, agglomerations of fluorescent cell fragments or debris may be mistaken for cells since only the total fluorescent area is considered instead of the size of individual objects.

The aim of the present work was to develop a software which can improve the field of automatic image analysis for zebrafish cancer models compared to previously published tools (Carreira et al., 2021; Yamamoto et al., 2019). Our approach processes three-dimensional confocal images and segments individual cells, which enables high-content image analysis for zebrafish larvae models of myeloid malignancies, as well as other cancers. The software was evaluated by investigating the proliferation of AML and MDS cells in zebrafish embryo and larvae, and their response to the leukaemia drugs DNR and Aza determined. We illustrate the use of single cell detection to extract additional information from confocal images, such as cell volume distributions and in vivo cell density maps. To evaluate the potential cardiotoxic effects of the administered treatments, we utilized a simple algorithm to automatically detect the larva heartrates from ten-second microscope videos.

## Results and discussion

### Cell segmentation

A large amount of data is contained in images obtained by confocal microscopy. However, this information can quickly be lost during data processing due to the feature extraction from three-dimensional confocal images being very time consuming and highly cumbersome. To alleviate this problem, we developed a software tool that automatically segments cells and further extracts relevant information from the acquired images. Here, we injected zebrafish larvae with fluorescent labelled cancer cells at two days post fertilization (dpf) and imaged daily using confocal microscopy until reaching 5 dpf. While Kimmel et al. defined the transition between the embryonic and larval stages to occur at after the protruding-mouth stage at 72h post fertilization, for convenience and to avoid confusion, we use the term zebrafish larva for all stages from the day of injection at 2 dpf (Kimmel et al., 1995).

An illustration of the cell segmentation is given in Figure 1. Figure 1-A to D give a montage of a zebrafish larva injected with MDS-L cells stained with the fluorescent marker CellTracker™ Deep-Red Dye and treated with daily azacitidine (Aza) injections over the course of four days. Iridophores are found on the dorsal region of the tail, characterized by dark spots on the brightfield image and slight signal in the far-red channel. Strong autofluorescence is also observed in the yolk sac, swim bladder and gastrointestinal tract, with strongest occurrence two days post injection (dpi), or 4 dpf, in Figure 1-C and G.

**Figure 1:**
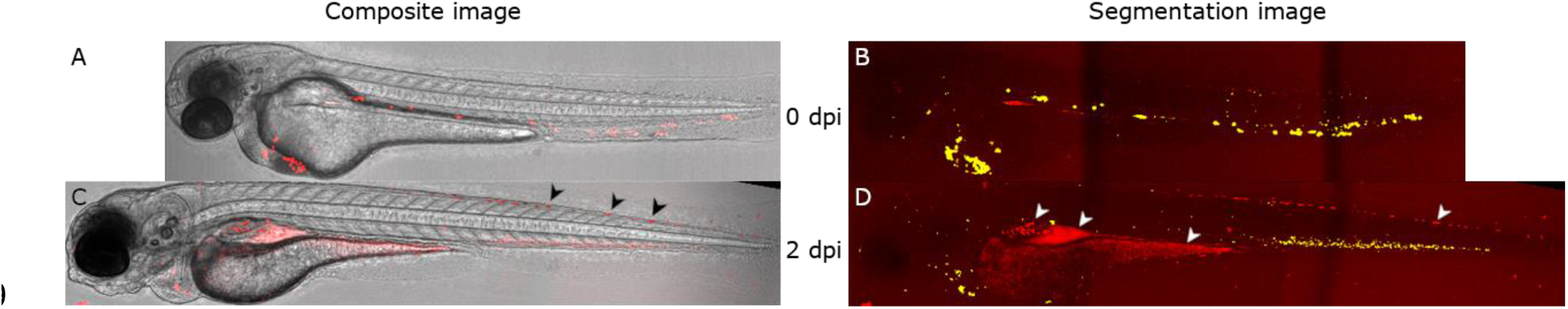
Segmentation of viable cells from confocal images of zebrafish larvae intravenously injected with the myelodysplastic syndrome cell line MDS-L. Approximately 4 nL of a 10 · 10^6^ cells/ml CellTracker™ Deep Red stained MDS-L cell suspension was injected into zebrafish larvae at 2 days post fertilization and imaged daily using confocal microscopy. The figure above illustrates one larva at the day of injection (0 dpi, A and B) and two days post injection (2 dpi, C and D). For illustration purposes, the confocal brightfield channel was sharpened using stack focusing in ImageJ (A and C), while the far-red fluorescence channel was projected using a max projection (B and D). Using our self-written ImageJ plugin, cells were segmented from the confocal images. Detected objects are illustrated with a green overlay, resulting in yellow where the red and green channels overlap (B and D). Sources of autofluorescence are shown in the larva at 2 dpi. In the composite image (C), Iridophores are indicated by black arrows, while in the segmentation image (D), the gut, yolk sack and iridophores are indicated by white arrows from left to right.

Figure 1-E to H visually illustrates the results of the cell segmentation performed by our software tool on the same image sets. Areas with high autofluorescence were manually masked prior to segmentation, resulting in the exclusion of iridophores, yolk sac, swim bladder and gastrointestinal tract from the set of detected objects. Automatic background correction can be achieved using different applications, such as the iterative threshold approach used by Carreira *et.al*. (Carreira et al., 2021). However, such approach did not yield satisfactory results in our data due to excessive removal of weakly fluorescent cancer cells, as well as inclusion of regions with high autofluorescence. Hence, a manual approach was chosen. A well-suited approach for removing background fluorescence would be fluorescent lifetime imaging. Where such a technique is not available, computational methods must be utilized. Automatic removal of background fluorescence using computational methods could be performed using machine learning or by including additional factors, such as location within the image or a comparison of multiple image channels.

One clear benefit of cell segmentation is that it collects additional data such as fluorescent intensity, volume, and precise position within the zebrafish larva. For instance, when transplanting cancer cells stained prior to injection, information of fluorescent intensity could be used to monitor proliferation of the cancer cells, since the fluorophore concentration in cells is halved for every division. This is important to be able to distinguish between treatments that kill cancer cells and those which for instance lead to senescence, as the latter would retain a higher fluorescent intensity compared to dividing cells. A quick illustration of plots comparing cell volumes to the average fluorescence of cells is given in Figure S1 for Molm-13 cells in medium, and Figure S2 and S3 for Molm-13 cells in zebrafish larvae with and without treatment, respectively. Additionally, since the boundaries and positions of cells are known, cut-outs containing single cells can be extracted from the original confocal image and further analysed in a manner similar to imaging flow cytometry.

### Identification of viable cancer cells based on volume distributions

When imaging fluorescently stained cells in zebrafish larvae, not every fluorescent object may represent a cell. Apoptotic bodies or cell fragment can still retain the fluorescent stain and be misidentified as viable cells. A method for separating these objects from viable cells can be achieved by using volume information to distinguish larger cells from smaller fragments.

To determine the lower threshold volume for viable cells, a volume distribution was created using the segmented objects of all injected embryos at the day of cell injection. As a comparison to cells in zebrafish larvae, cells stained with CellTracker™ Deep-Red were kept in medium and imaged using confocal microscopy. The resulting confocal image and image illustrating the segmentation results of Molm-13 cells are shown in Figure 2-A on the top and bottom, respectively, with the resulting volume distribution is shown in Figure 2-B. As the Molm-13 and MDS-L cells have an approximately equal size, this control is only shown for Molm-13.

**Figure 2:**
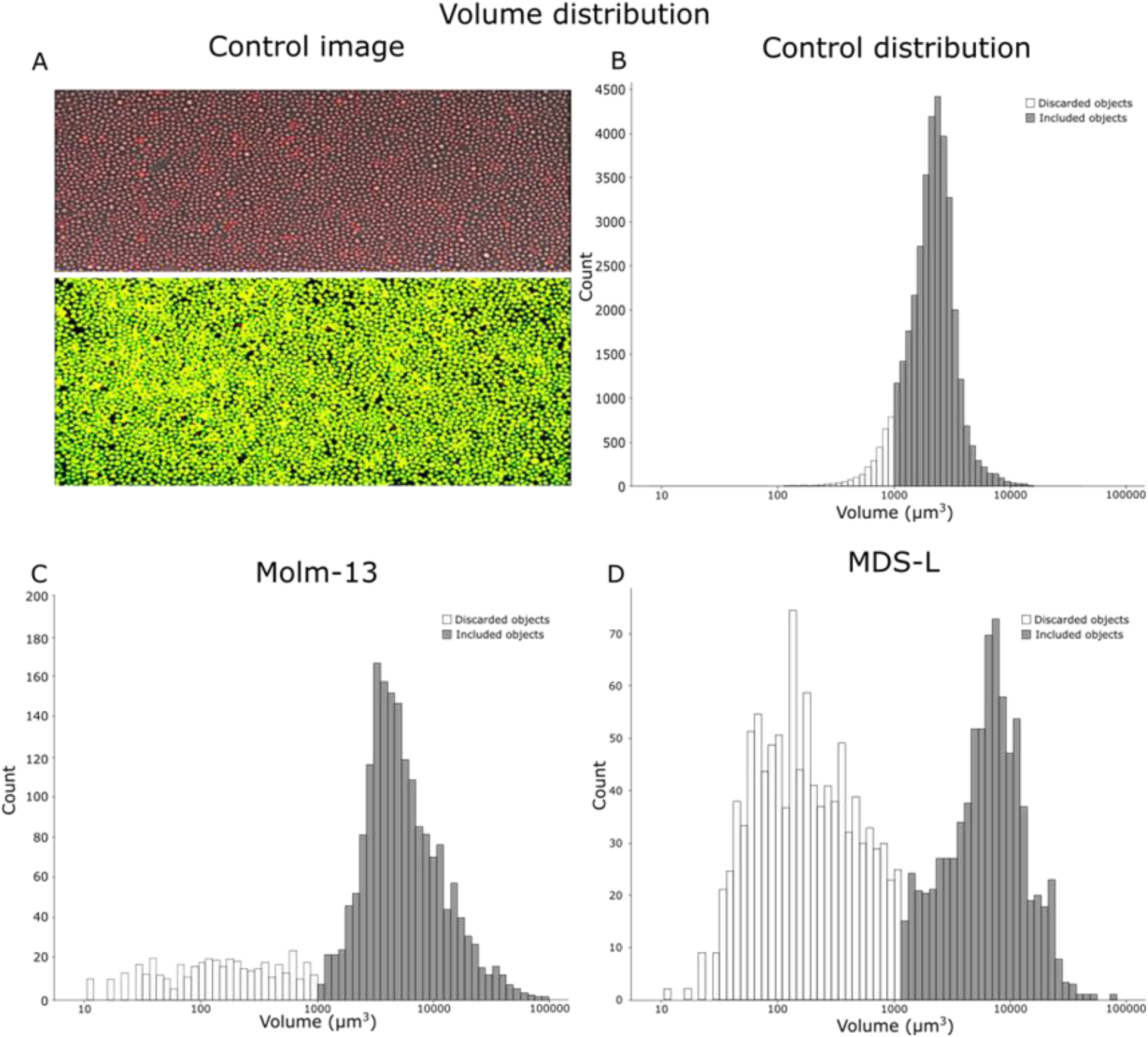
Measurement of injected cancer cell volume distributions based on confocal images. A and B: Cultured Molm-13 AML cells were stained with CellTracker™ Deep Red and imaged by confocal microscopy. An illustration of the segmentation process is shown in A, with the composite fluorescence and brightfield image shown at the top and the resulting segmentation below (composite image of red fluorescence and segmented overlay in green). B shows the cumulative volume distribution of all segmented cells. C and D: For volume distributions of cancer cells in zebrafish larvae, 4 nL of 10 · 10^6^ cells/ml CellTracker™ Deep Red stained cancer cell suspensions were injected into the posterior cardinal vein of 18 zebrafish embryos at 2 days post fertilization. Following cell injection, the embryos were imaged by confocal microscopy, and the images processed as the control samples in A and B. Volume distributions for the cell lines Molm-13 and MDS-L in zebrafish larvae are given in C and D, respectively. Using the volume distribution from B, a lower volume cut-off for viable cells was determined. The cell populations above and below this threshold are shown in grey and white, respectively.

The volume distributions for Molm-13 and MDS-L in zebrafish larvae at the day of injection are shown in Figure 2-C and D, respectively. Based on these volume distribution plots, the volume thresholds for viable cells were determined to be above ∼1000 μm^3^ for both Molm-13 and MDS-L. This volume responds to an equivalent spherical diameter of above 12 μm. The measured size can however be impacted by scattering in the z-plane when acquiring confocal images. This will cause objects to appear stretched along the z-axis and can impact accurate measurement of volumes. However, all measured objects are affected in a similar manner, and their relative differences are still readily detectable. Compared to Molm-13, the size distribution of MDS-L exhibits two populations at the day of injection (Figure 2-A and B). The population consisting of objects with smaller volume as shown in white is likely to be debris originating from dead cells that did not tolerate the handling during staining and injection prior to imaging.

### Distribution of injected Molm-13 and MDS-L cell lines in zebrafish embryos

The position of each detected object with a volume above the predetermined threshold of 1000 μm^3^ was recorded during cell segmentation. Using this data, density maps of injected Molm-13 or MDS-L cells in zebrafish larvae were constructed as seen in Figure 3 and Figure 4, respectively. It is important to note that the density maps are relative to themselves. This makes each distribution clearer; however, changes in cell counts between density maps are no longer reflected.

**Figure 3:**
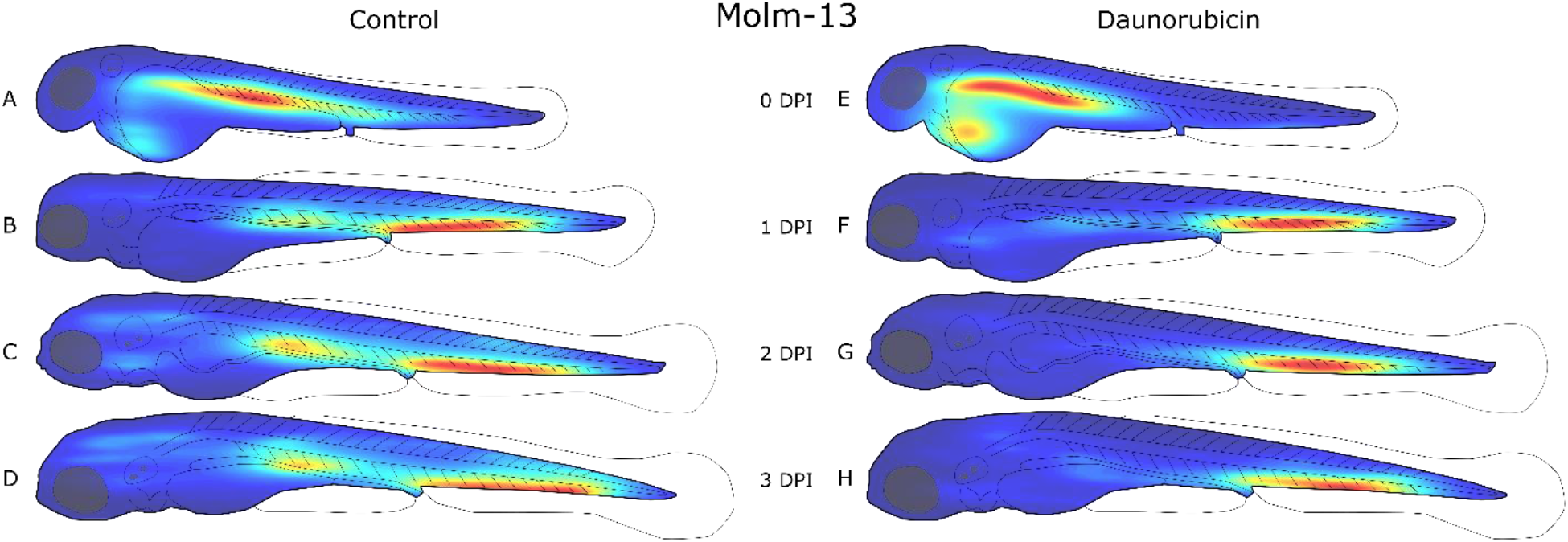
Distribution of Molm-13 cells in zebrafish larvae after intravenous injection with daunorubicin. Zebrafish embryos were injected with approximately 4 nL of a 10 · 10^6^ cells/ml CellTracker™ Deep Red stained Molm-13 cell suspension at 2 days post fertilization (dpf), and either left untreated (A-D) or treated with a 4 nL injection of 1mM daunorubicin (E-H). Each larva was imaged daily using confocal microscopy, and the images analysed using the self-written ImageJ plugin as described in the method section. The location of each cell above the volume threshold determined in Figure 2-C was used to create a distribution map for each group on each day post injection (dpi). The density map was generated using MATLAB and visualizes the areas of highest cell density (red) and lowest density (blue). Each density map is relative to itself. This results in clearer distributions, however changes in cell counts between density maps are no longer reflected.

**Figure 4:**
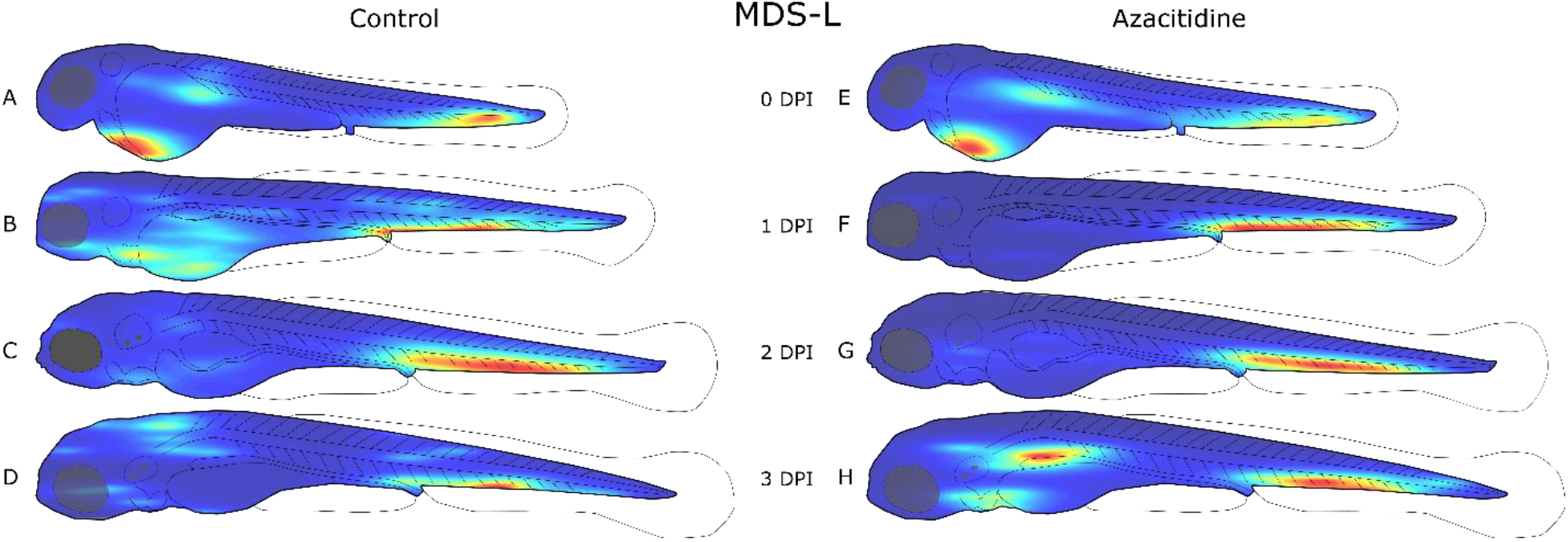
Distribution of MDS-L cells in zebrafish larvae after intravenous injection with azacitidine. Zebrafish embryos were injected with approximately 4 nL of a 10 · 10^6^ cells/ml CellTracker Deep Red stained MDS-L cell suspension was injected into 18 zebrafish embryos at 2 days post fertilization (dpf), and either left untreated (A-D) or treated with daily 4 nL injections of 1mM azacitidine (E-H). Each larva was imaged daily using confocal microscopy, and the images analysed using the self-written ImageJ plugin as described in the method section. The location of each cell above the volume threshold determined in Figure 2-D was used to create a distribution map for each group on each day post injection (dpi). The density map was generated using MATLAB and visualizes the areas of highest cell density (red) and lowest density (blue). Each density map is relative to itself. This results in clearer distributions, however changes in cell counts between density maps are no longer reflected.

At the day of injection, a high cell density can also be seen around the injection site in the posterior cardinal vein in Molm-13 (Figure 3-A and E), and in the heart region for MDS-L (Figure 4-A and E). From the day after injection (1 dpi) until the end of the experiments (2 dpi), both cell lines were found to aggregate in the ventral tail region around the caudal vein. This region contains the caudal haematopoietic tissue from two to five dpf (Gore et al., 2018). A possible explanation for this accumulation could be a selective exit from the bloodstream by the cancer cells at a haematopoietic niche due to cytokines which also attract human leukaemia cells, whose natural environment is the bone marrow.

The Molm-13 cell distributions obtained from larvae treated with DNR (Figure 3-E to H) show a lower cell density around the posterior cardinal vein compared to the untreated transplanted control, while the population in the ventral tail around the caudal vein remains strong. The reduction of cancer cells around the posterior cardinal vein can be a result of the injection site of DNR being in the same area; however, the high injection velocity of the drug, as well as functioning circulatory system likely result in a rapid and even distribution of the drug in the blood. Thus, a more likely explanation is that the AML cells are more protected from DNR when residing in the haematopoietic niche ventral to the caudal vein.

In larvae injected with MDS-L, the regions outside of the haematopoietic niche were found to have a higher cell density in control larvae without injection compared to larvae treated with Aza (Figure 4 A-D and E-H, respectively). A notable outlier can be observed in Figure 4-H, where a high cell density can be found around the posterior cardinal vein in Aza-treated larvae when compared to control larvae (Figure 4-D). The MDS-L cells are therapy resistant, and the new population around the posterior cardinal vein could be the formation of a new colony outside the caudal haematopoietic tissue.

### Quantification of tumour burden in zebrafish larvae

As well as determining the positions of segmented cells, the software tool also measures cell count and volumes. Using this information, the temporal development of tumour burden was compared in larvae between untreated larvae and larvae receiving treatment. We observed a decline in tumour burden in untreated larvae throughout the three-day observation period (Fig. 5). The reason for this decline can be due to the stress cells are subjected to from handling before and during the injection, as well as the temperature change from 37 °C in the incubator to 31 °C in the zebrafish. This initial rapid decline during the first 24 hours is followed by less dramatic decrease at 2 and 3 dpi for Molm-13 (Figure 5-A and B). This can be an indication that the cancer cells now have adapted to the new microenvironment. Such an initial decline in engrafted cells is also seen in mammalian cancer models such as mice, where patient derived xenografts in immunosuppressed mice have an initial latency period followed by proliferation (Siolas and Hannon, 2013).

**Figure 5:**
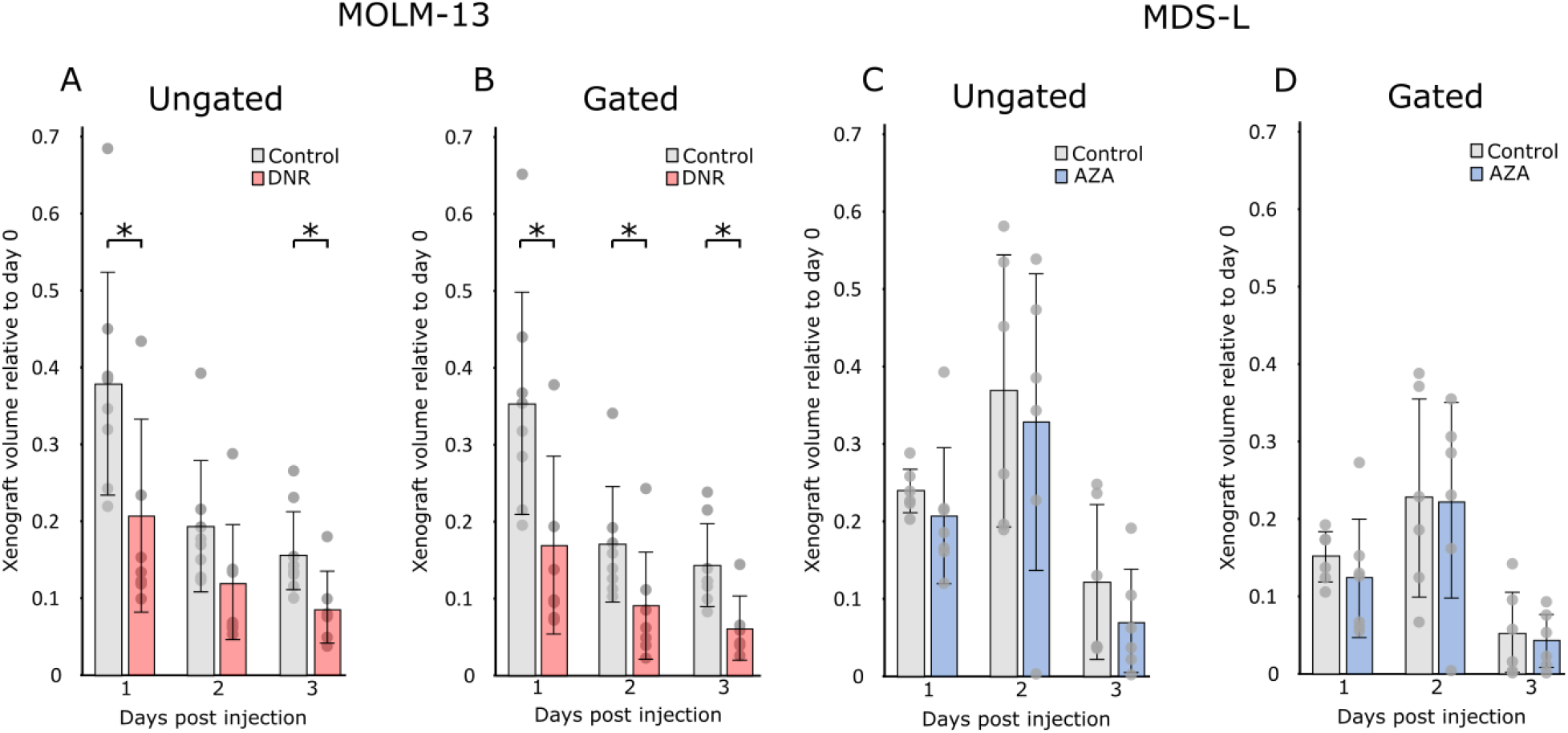
Development of tumour burden of Molm-13 and MDS-L in zebrafish larvae. Molm-13 and MDS-L cells were stained with CellTracker™ Deep Red and 4 nL of a 10 · 10^6^ cells/ml suspension engrafted into zebrafish larvae two days post fertilization by injection into the posterior cardinal vein. Following engraftment, the larvae were daily using spinning disk confocal microscopy. The treatments consisted of a single 4 nL 1mM daunorubicin injection at the day of the transplant for Molm-13 cells-xenografted larvae, whereas for larvae engrafted with MDS-L, treatment consisted of daily injections of 4 nL 1mM azacitidine. Imaging and treatment were continued until the embryos reached 5 days post fertilization. Using our plugin, cells were counted and segmented from the confocal images. The total volume of all segmented objects in larvae engrafted with Molm-13 and MDS-L are shown in A and C, respectively. Using the volume threshold of 1000 µm^3^ determined from the images in Figure 2 C and D, the filtered total cell volume was determined (B and D). Significance between treatments and controls were found using two-tailed Welch’s t-test. *:P<0.05

For Molm-13, the summed volumes without and with volume gating are shown in Figure 5-A and B, respectively. As seen in Figure 5-B, larvae treated with DNR (red) displayed a significantly lower tumour burden compared to the untreated (grey) throughout the observation period. An impact on the volume distribution of Molm-13 cells due to the DNR treatment can also be seen as shown in Figure S4. The cell volume sum of MDS-L cells without and with volume gating are shown in Figure 5-C and D, respectively. A large difference between the gated and non-gated cell volumes can be seen. This can reflect the population of smaller cell fragments that were also seen in the volume distribution in Figure 2-D. In contrast to Molm-13 cells treated with DNR, no significant difference in total tumour burden can be seen in MDS-L cells treated with Aza. This can be attributed to the slow mechanism of action for hypomethylating drugs as observed in in vitro studies, but also that the MDS-L cell line is therapy resistant, and not likely to respond to this drug.

### Monitoring cardiotoxic effect from anti-cancer drugs Daunorubicin and Azacitidine

The use of zebrafish larva as a model for cardiotoxic drugs has been demonstrated in several studies (Maciag et al., 2022; Zhu et al., 2014). Usually, the cardiotoxic effects are monitored by manually counting the heartrate. In addition to being time-consuming, this can also lead to biased results if not performed blindly. However, by using a software to count the heartrate of zebrafish embryo, both these issues are eliminated. The macro used for these measurements uses intensity fluctuations in the acquired microscope videos due to the larva’s heartbeat. A detailed description of this is given in the methods section.

Previous studies have shown that anthracyclines reduce heart rate in zebrafish larvae (Han et al., 2015), and we wanted to test whether we could observe this effect using our macro for calculating heart rate. A reduction in heartrate for DNR-treated larvae occurred only at one dpi, but not at two and three dpi (Figure 6-D). This could be because of elimination of DNR by metabolism or excretion. Also, zebrafish are known to be able to regenerate tissue, and heart regeneration has been observed in adult fish (Poss et al., 2002).

**Figure 6:**
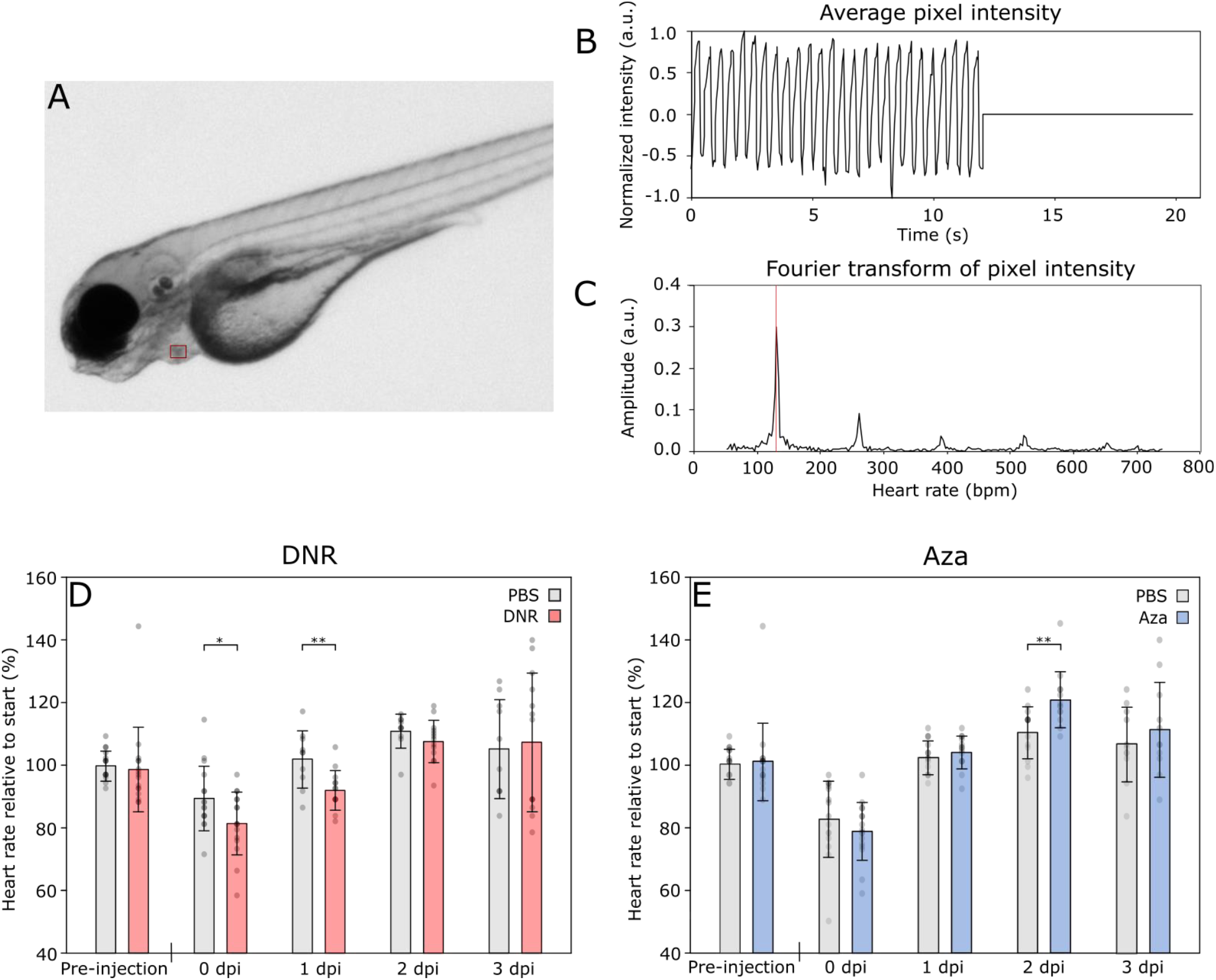
Aza and DNR effects on zebrafish heart rate. The zebrafish larvae heart rate in the cardiotoxicity assays was found by filming the zebrafish for 12 seconds and using a self-written ImageJ macro to calculate the heart rate from the obtained video as described in the methods section. A) Still image from a film analysing zebrafish larvae heart rate. The red rectangle marking the selected area (SA) for pixel intensity measurement. B) Average pixel intensity in the SA. C) Fourier transform of pixel intensity from the SA to calculate beats per minute (BPM). D and E) Heart rate of zebrafish larvae injected with 4 nL PBS, Aza (1 mM, D) or DNR (1 mM, E). For the Aza-test, the zebrafish larvae were injected daily after Pre-injection, 1 dpi and 2 dpi measurements. For the DNR-test, the zebrafish larvae were injected once, after the Pre-injection observation. The heart rate was related to the average heart rate of the control group pre-injection. Significance found using two-tailed Welch’s t-test. *: P<0.05, **: P<0.01.

In the fish treated with Aza, there was an increased heartrate at two dpi (Figure 6-E). The effect could be a stress reaction of multiple injections of the cytostatic drug. In contrast to DNR, the effect of Aza on the heart of zebrafish is little studied, but one report showed reduced survival, malformations and cardiac effects such as pericardial oedema and reduced ventricular volume at 72 hpf when injected with the drug at the one to four cell stage (Yang et al., 2019). In humans, cases of pericardial effusion and pericarditis have been reported as a result of Aza treatment (Goo et al., 2019; Newman et al., 2016). In our tests, pericardial effusion as an oedema surrounding the heart was observed in two out of twelve of the Aza recipients at three dpi, but none of the recipients of PBS injections (Figure S5). This could indicate that the heart of the zebrafish larvae responds to Aza in a similar manner as the human heart.

In this study we performed intravenous injections of DNR and Aza. Administration of drugs through the embryo water is a more frequently applied method, and while administration through water could give a much more predictable and stable drug concentration, it is not always known to which extent the different drugs are absorbed through the skin of the zebrafish. Moreover, Aza is highly unstable in aqueous solutions and for our purpose not a viable route of administration.

## Conclusion

In summary, we presented a software tool capable of extracting a large range of information on a single cell basis from confocal images of zebrafish larvae injected with fluorescently stained cancer cells. While zebrafish larvae were used as subjects in this project, this software is readily adaptable for any confocal image containing fluorescently labelled cells. To facilitate the monitoring of heart function in intact zebrafish larvae, we developed a simple procedure to measure larval heartrates. Future work on reliably and automatically detecting the outline of zebrafish larvae in confocal images as well as removing background fluorescence can eliminate the need for user interaction entirely and further speed up the acquisition of high-content data from confocal images.

## Materials and Method

### Materials

Microinjection pipettes for cell transplantation (BioMedical Instruments, VESbv-11-0-0-55), microinjection pipettes for drug injection (BioMedical Instruments, VICbl-4-0-0-55), Daunorubicin (DNR) (Cerubidine, Sanofi-Aventis), Azacitidine (Aza) (Sigma-Aldrich, A2385-100MG), RPMI-1640 Medium (Sigma-Aldrich, R5886-6×500ML), Ethyl 3-aminobenzoate methanesulfonate (Tricaine) (Merck Life Sciences, E10521-50G), Foetal bovine serum (FBS)(Sigma-Aldrich, F7524-500ML), L-glutamine (Sigma-Aldrich, G7513-100ML), Penicillin-Streptomycin (Sigma-Aldrich, P0781-100ML), Recombinant Human IL-3 Protein (IL-3) (BioTechne, 203-IL-010), CellTracker™ Deep Red Dye (ThermoFisher, C34565)

### Methods

#### Zebrafish maintenance

The transparent zebrafish line Casper was used (White et al., 2008). Fertilized zebrafish eggs were obtained from the zebrafish facility at the University of Bergen. This facility is run according to the European Convention for the Protection of Vertebrate Animals used for Experimental and Other Scientific Purposes. The zebrafish eggs, embryos and larvae were kept in petri dishes with E3 medium (5 mM NaCl, 0.17 mM KCl and 0.33 mM MgSO4) with added methyl blue at 28.5 °C. Petri dishes were cleaned daily for debris or dead embryos/larvae. Following injection of cancer cells, the zebrafish larvae were kept at 31 °C as a compromise between the 28.5 °C preferred by the larvae and 37 °C the injected cells. All zebrafish larvae were euthanized at five dpf. For euthanasia, the zebrafish larvae were first kept on ice for at least 20 minutes before being frozen at -20 °C over-night. Due to no zebrafish larvae exceeding five dpf, no application to the ethics committee was required.

#### Cell line maintenance

The AML cell line MOLM-13 and MDS cell line MDS-L were cultured at 37 °C in a 5% CO2 atmosphere in RPMI medium (Matsuo et al., 1997; Rhyasen et al., 2014). The medium was enriched with 10% FBS, 20 mM L-glutamine, 100 IU/ml penicillin and 0.1 mg/ml streptomycin. For the cell line MDS-L, the medium was additionally enriched with IL-3 to a concentration of 25 ng/ml. Both cell lines were kept at a concentration between 100.000 and 1.000.000 cells /ml during routine maintenance.

#### In-vivo cell experiments

##### Cell staining

The cells were stained using CellTracker™ Deep Red Dye following the manufacturer’s protocol. In brief, cells were centrifuged at 90 RCF for 5 minutes, the medium decanted, and the cells resuspended in fresh serum free RPMI medium with 20 μM CellTracker™ staining solution. The cells were incubated for 30 minutes at 37 °C before being centrifuged at 90 RCF for 5 minutes and resuspended in fresh medium to a concentration of approximately 10 million cells/ml for injection into zebrafish larvae.

##### Transplantation of human leukaemia cells in zebrafish embryo

Zebrafish larvae at 2 dpf in the long-pec stage according to the developmental stages proposed by Kimmel *et.al*., were anesthetised using a 0.7 mM tricaine solution and mechanically dechorionated under a microscope using forceps (Kimmel et al., 1995). A microinjection pipette with an inner diameter of 11 µm was filled with CellTracker™ Deep-Red stained cell suspension and mounted onto a Narishige MMN-5 with MMO-220A micromanipulator system and connected to an Eppendorf FemtoJet 4x microinjector. The injection pressure and time were adjusted to achieve an injection volume of 4 nL. The anesthetised zebrafish larvae were placed on a 2% agarose bed and injected into the posterior cardinal vain as illustrated in Figure 7. Following injection, the zebrafish were kept anesthetized and transferred to an 18 well confocal slide. For imaging, an Andor Dragonfly 505 confocal microscope was used. The brightfield channel was used to visualize the larva, while a 700/38nm band-pass filter with a 637nm excitation laser was used to track the injected cancer cells.

**Figure 7:**
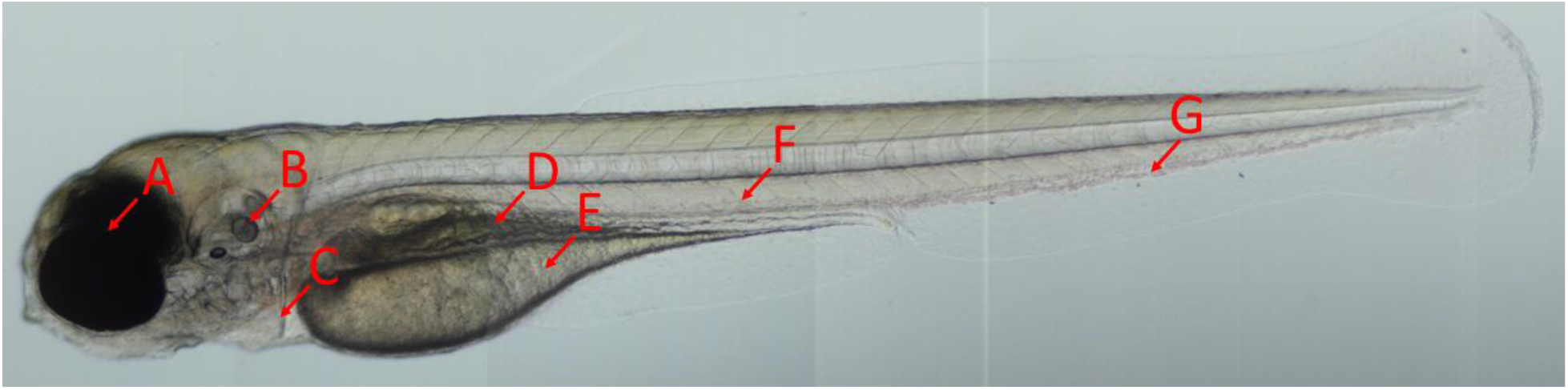
Overview of a zebrafish embryo. A zebrafish larva imaged at four days post fertilization using bright field microscopy. The arrows indicate the eyes (A), otolith (B), heart (C), gut (D) and yolk sac (E). Intravenous injections were performed in the posterior cardinal vein (F) down-stream from the caudal vein (G).

After confocal imaging, half of the larvae were additionally injected into the posterior cardinal vein with an anti-cancer drug, using the same protocol as for cell injections apart from using an injection pipette with a smaller inner diameter of 4 µm. DNR injections were only performed once on the same day as cell injections, while Aza was injected daily after each confocal imaging due to the molecule’s short half-life in aqueous solutions

The following days, the zebrafish embryos were imaged daily using confocal microscopy. Prior to reaching 120 hours post fertilization (5 dpf), all embryos were euthanized. The acquired confocal images were analysed using the self-written software tool.

#### Software tool for image processing

A plugin for the image processing software ImageJ (Version 1.53c, (Schneider et al., 2012)) was written in Java using JDK 1.8. The plugin was written using the integrated development environment Apache NetBeans IDE 13. A graphical overview of the software tool’s workflow is given in Figure 8.

**Figure 8:**
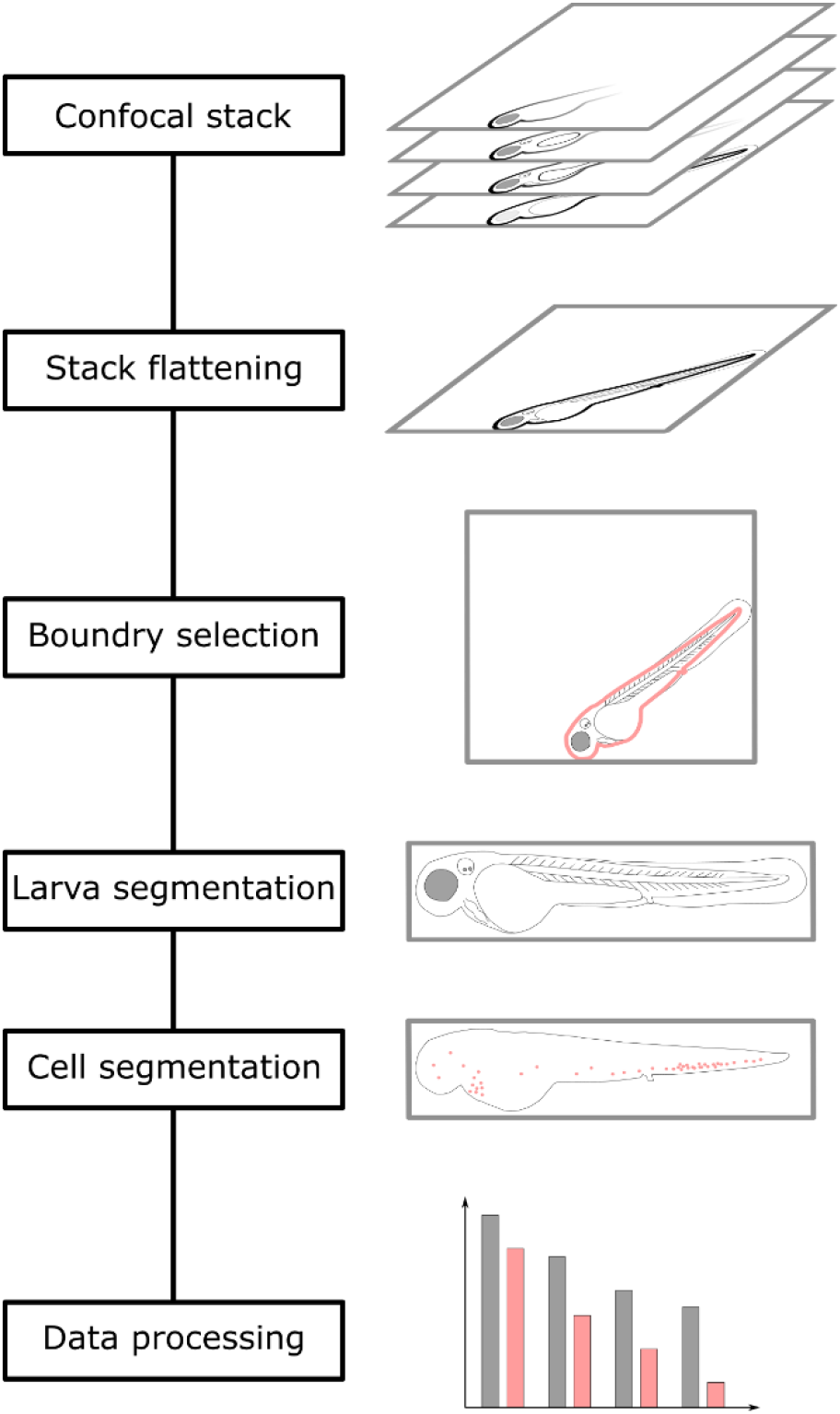
Workflow of the zebrafish analysis tool. The acquired confocal files are first flattened to a 2D representation to enable easy visual analysis. Following flattening, the larval boundaries are selected by the user to determine the location and orientation of the embryo. Using this information, the larvae are segmented and realigned to a standardized orientation. Background levels are determined by the user and all objects located within the larva segmented using a watershed algorithm. Following these steps, the data can be exported for further analysis.

The software starts by flattening each confocal stack to a 2D representation. Importing the confocal image files is enabled using the Bio-Formats library (Linkert et al., 2010). The brightfield channels are flattened by utilizing code from the Stack-Focuser plugin originally written by Michael Umorin, but slightly modified to suit our needs (https://imagej.nih.gov/ij/plugins/stack-focuser.html). Fluorescent channels are flattened using a max projection. Following flattening, the borders of the embryos are manually selected by the user. Using the selected outline, the angle of the larva is determined by the selection boundary’s Feret angle, while position of the yolk sac is determined using a distance map by identifying the furthest internal point from the boundary. Using the position and orientation of each embryo, a montage can automatically be created for each larva from the day of injection to the day of euthanasia.

For cell segmentation, the level of background fluorescence is determined manually by the user, as well as masking of any sources of high autofluorescence, such as iridophores, the gastrointestinal tract and yolk sac. Cell segmentation is performed using a watershed algorithm found in the mcib3d library, with the code slightly modified to increase segmentation speed (Ollion et al., 2013). To find starting points for the watershed algorithm, the fluorescent stack is first pre-processed using a 3D-median filter with radius 3 for x and y, and 2 for z. This is followed by a local maximum filter with a cut-off value of the predetermined background fluorescence. Using the obtained seed image as a starting point, a watershed is performed on the fluorescent stack. This process is repeated for each larva for each daily acquisition, and the detected objects stored.

Following cell segmentation, a size distribution of each detected object is created. From this distribution, a size range for viable cells is determined. Using this size range as a filter, the cell count and total cell volume in each larva is exported for further analysis.

From the location-data obtained from the segmentation process, a heatmap is constructed to visualize cell distributions. X- and Y-coordinates of segmented cells for all larva within the same group and day are cumulated and a heatmap generated using MATLAB (R2021 Update 2) with a modified version of the Data Density Plot plugin supplied by Malcolm McLean (https://www.mathworks.com/matlabcentral/fileexchange/31726-data-density-plot). All heatmaps are normalized to facilitate the visualization of the cell distribution.

#### Cardiotoxic assay

Two dpf zebrafish larvae were intravenously injected with the drugs. To ensure that the effect of temperature and tricaine was similar in all larvae, the zebrafish larvae were exposed to 0.7 mM tricaine and left in room temperature for at least 30 minutes prior to filming. The zebrafish larvae then were injected with either of the following into the PCV: 4 nL DNR (1 mM), Aza (1 mM) or PBS as control. Due to the rapid decomposition of Aza, the injection of this drug and PBS for the representative control group was repeated at three and four dpf. DNR and its PBS control group was only injected at two dpf.

To determine heart rate, each larva was filmed for 12 seconds using a Leica M205 stereo microscope fitted with Leica DFC3000 G camera and the Leica Application Suite X software. At two dpf, the zebrafish larvae were filmed both before (Pre-injection) and directly after (0 dpi) the injection. This was to observe whether there were immediate toxic effects of the drugs. For simplicity, the time points are related to the first injection for all groups.

The heart rate was found using a self-written macro for FIJI. To determine the heartrate, a region around the heart is selected. The macro measures the average intensity in each frame of the video and normalizes the values to a range between -1 and 1. To comply with the requirements of the fast Fourier transform (FFT) algorithm utilized by FIJI, artificial measurements with the value zero are added to the measurement list until the total number of measurements equals a power of two. After performing the FFT, the heartrate is determined to be the fundamental frequency of the frequency plot.

#### Statistical analyses

The data in all bar charts is presented as average with standard deviation. Statistical significance was determined using a two-tailed Welch’s t-test performed using RStudio for Windows, version 2022.02.2 Build 485 (RStudio, PCB, Boston, MA, USA).

## Acknowledgements

We thank the Molecular Imaging Centre (MIC) at the University of bergen for support and training on confocal microscopy, as well as the zebrafish facility at the University of bergen for access to E3 medium and mature zebrafish for breeding.

## Competing interests

The authors declare that they have no competing interests.

## Funding

This project was funded by the Western Norway Health Authorities (grant no: F-12533) and Nordforsk (the NordForsk Nordic Center of Excellence “NordAqua” (No. 82845), and the Norwegian Society for Children’s Cancer.

## Supplementary information

**Figure S1.**
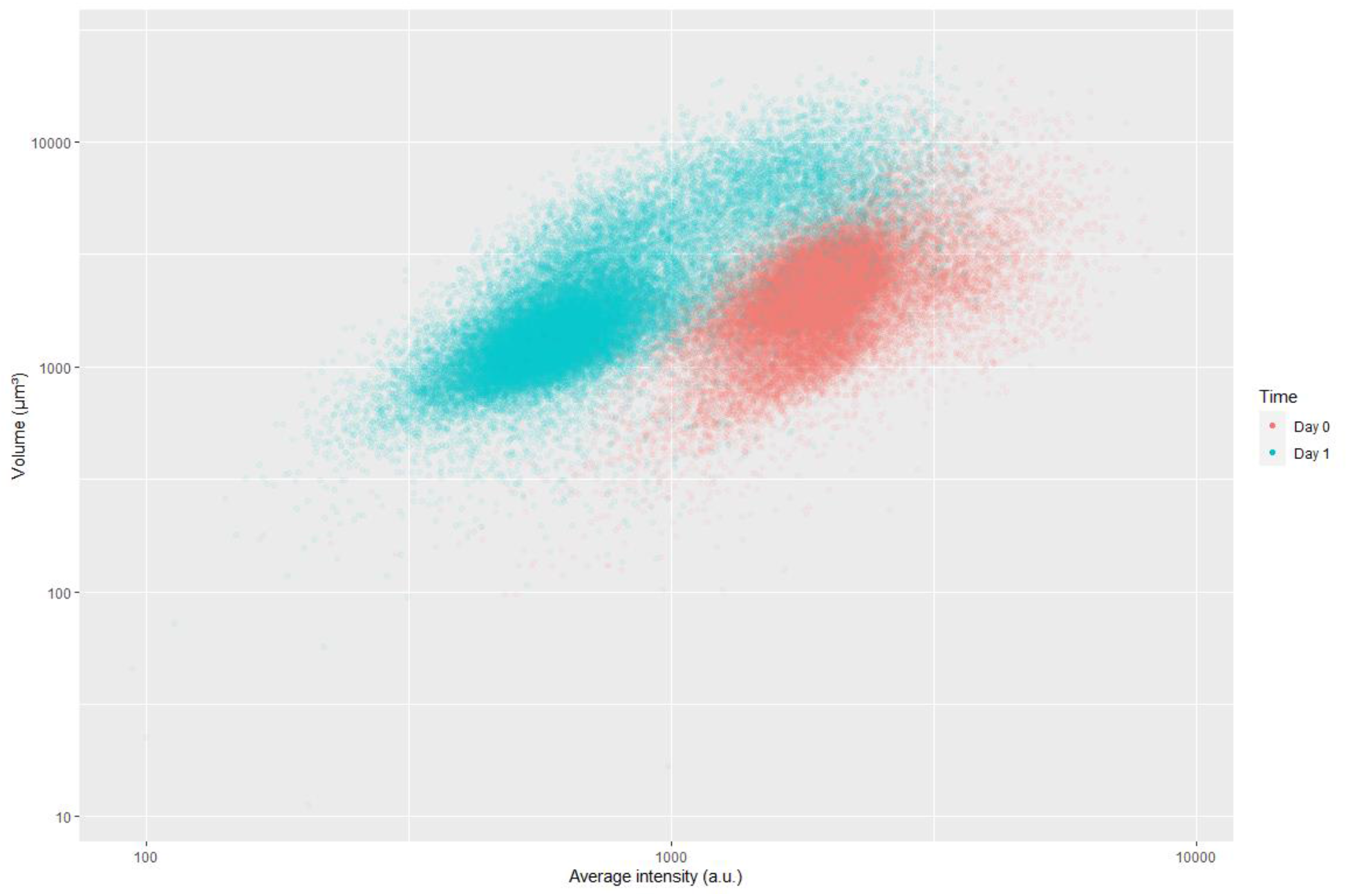
Scatter plot of intensity vs volume of Molm-13 cells in medium from confocal images

**Figure S2.**
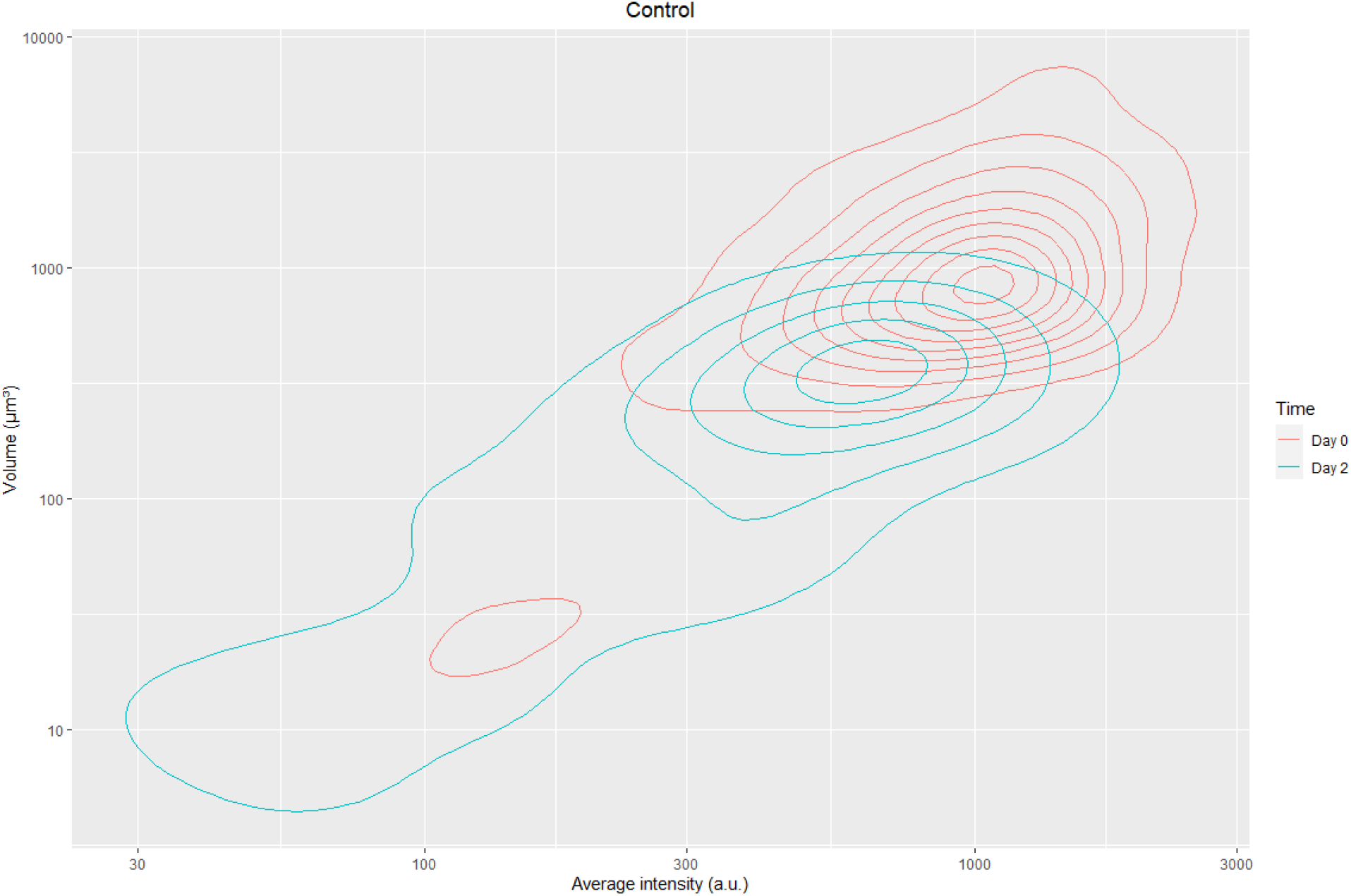
Density plot of fluorescent intensity vs volume of molm-13 cells transplanted in zebrafish without treatment.

**Figure S3.**
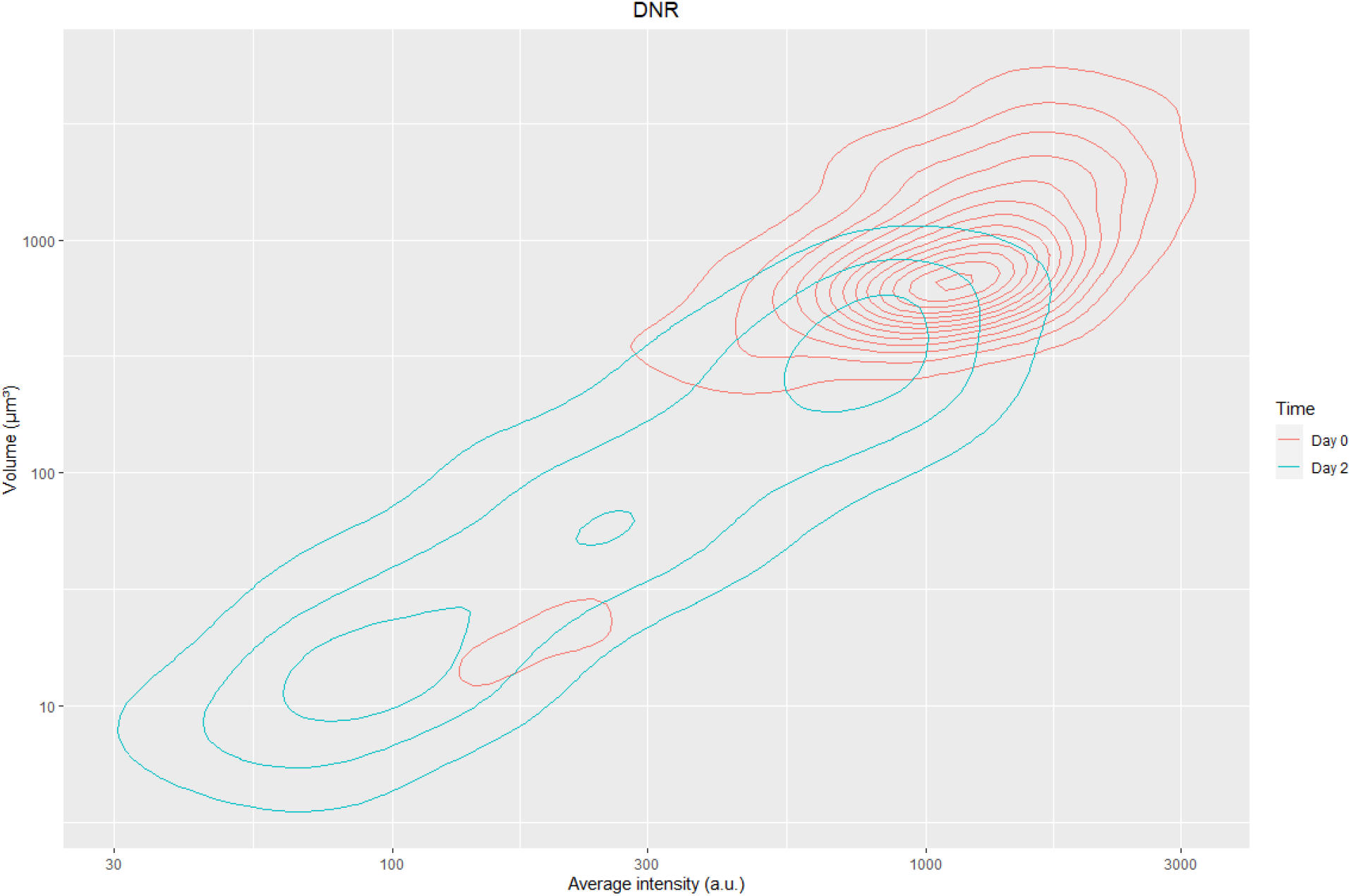
Density plot of fluorescent intensity vs volume of molm-13 cells transplanted in zebrafish with daunorubicin treatment.

**Figure S4.**
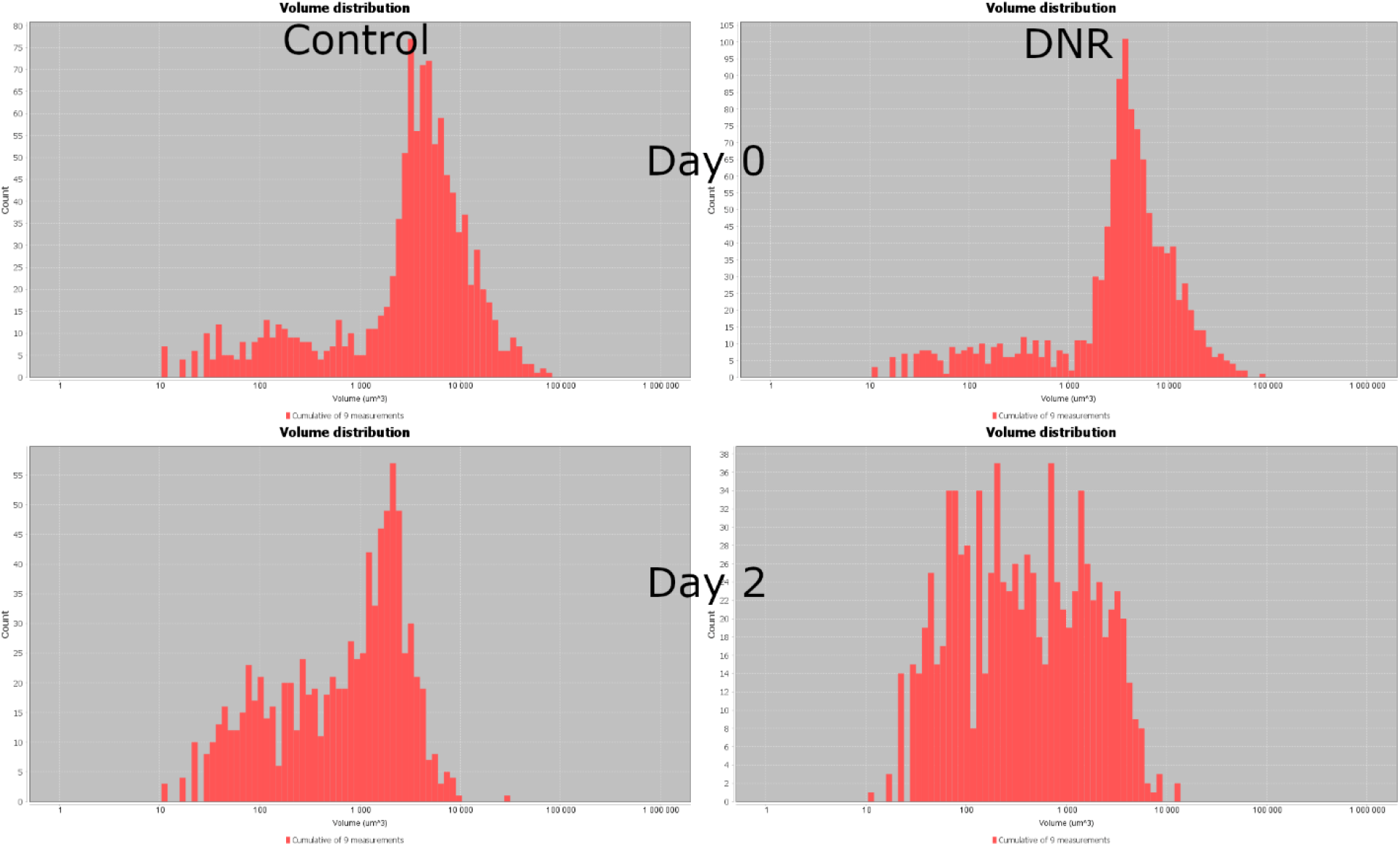
Volume distribution of Molm-13 cells transplanted in zebrafish larvae

**Figure S5.**
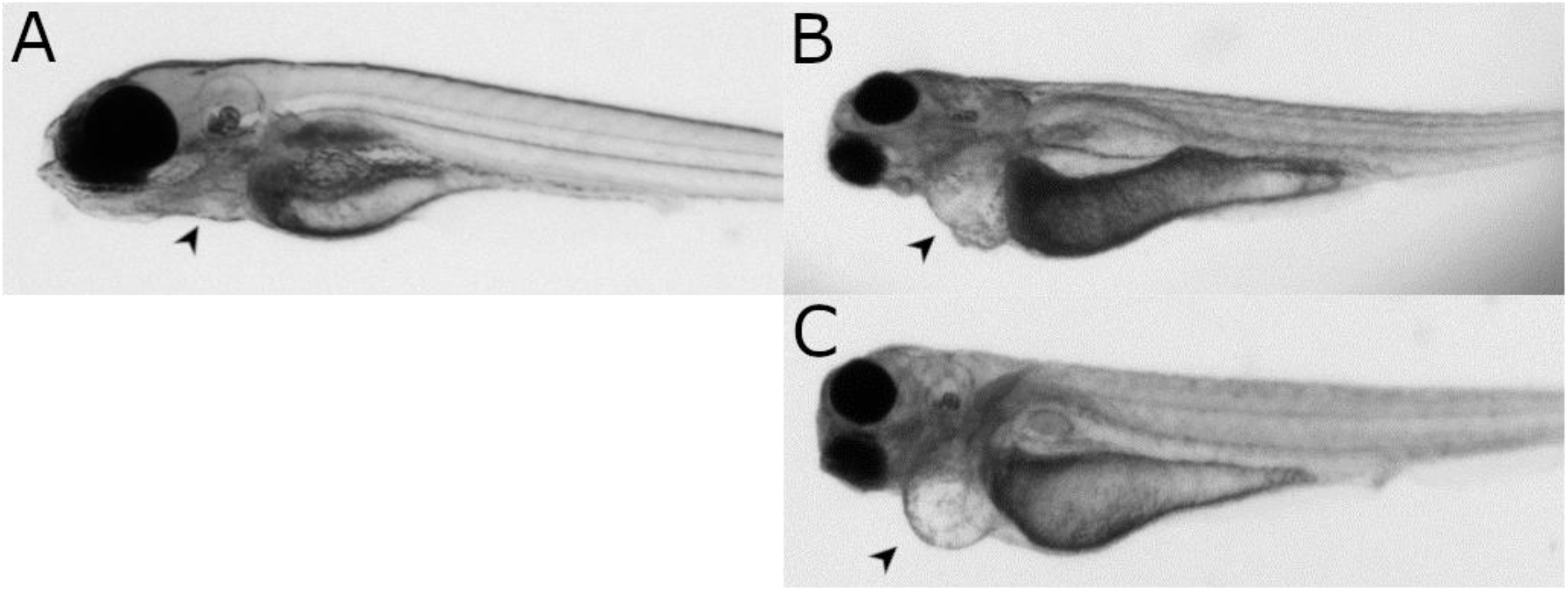
Untreated (A) and Aza-treated zebrafish larvae (B and C). Note the presence of mild and severe oedema in B and C, respectively.

## Notes

### Competing Interest Statement

The authors have declared no competing interest.

